# The nucleocapsid (N) proteins of different human coronaviruses demonstrate a variable capacity to induce the formation of cytoplasmic condensates

**DOI:** 10.1101/2024.11.20.624003

**Authors:** Maria A. Tikhomirova, Oleg L. Kuzmenko, Eugene A. Arifulin, Olesya M. Shirokova, Yana R. Musinova, Eugene V. Sheval

## Abstract

To date, seven human coronaviruses (HCoVs) have been identified. Four of these viruses typically manifest as a mild respiratory disease, whereas the remaining three can cause severe conditions that often result in death. The reasons for these differences remain poorly understood, but may be related to the properties of individual viral proteins. The nucleocapsid (N) protein plays a crucial role in the packaging of viral genomic RNA and the modification of host cells during infection, in part due to its capacity to form dynamic biological condensates via liquid-liquid phase separation (LLPS). In this study, we investigated the capacity of N proteins derived from all HCoVs to form condensates when transiently expressed in cultured human cells. A fraction of the transfected cells were observed to contain cytoplasmic granules in which the most of the N proteins were accumulated. Notably, the N proteins of SARS-CoV and SARS-CoV-2 showed a significantly reduced tendency to form cytoplasmic condensates. The condensate formation was not a consequence of overexpression of N proteins, as is typical for LLPS-inducing proteins. These condensates contained components of stress granules (SGs), indicating that the expression of N proteins caused the formation of SGs, which integrate N proteins. Thus, the N proteins of two closely related viruses, SARS-CoV and SARS-CoV-2, have the capacity to antagonize SG induction and/or assembly, in contrast to all other known HCoVs.

## Introduction

The emergence of the novel human coronavirus (HCoV) SARS-CoV-2, which causes the respiratory disease known as COVID-19, has become a significant global public health concern, prompting extensive research into the biological characteristics of the virus and the disease it causes. SARS-CoV-2 is the seventh member of the Coronaviridae family known to infect humans (Su et al., 2016). The four HCoVs - HCoV-229E, HCoV-NL63, HCoV-HKU1, and HCoV-OC43 - typically cause mild to moderate upper respiratory tract diseases and are estimated to account for 15–30% of human colds (Liu et al., 2021a). Other HCoVs - SARS-CoV, MERS-CoV, and SARS-CoV-2 - are highly pathogenic and can infiltrate the lower respiratory tract, leading to severe respiratory failure and a high death rate.

Different strains of SARS-CoV-2 have been observed to demonstrate varying abilities to infect lung cells (Valyaeva et al., 2022). However, the processes occurring within infected cells also exert an influence on the severity of the disease. This duality is confirmed by genomic analysis which revealed that low pathogenic HCoVs diverge from highly pathogenic HCoVs in two proteins, S and N (Gussow et al., 2020). The S protein, also known as the spike protein, is essential for cell penetration, while the N protein, or nucleocapsid protein, plays a pivotal role in packaging and organizing genomic viral RNA in SARS-CoV-2 virions (Wu et al., 2023). Consequently, the characteristics of the N protein may significantly impact the severity of disease.

The amount of SARS-CoV-2 N protein depends on the stage of infection - it is low in the early stages, then increases dramatically as the infection progresses, resulting in N protein becoming the most abundant viral protein in infected cells (Kim et al., 2020, 2021; Grossegesse et al., 2022). N protein can interact with different cellular proteins (Gordon et al., 2020; Chen et al., 2021; Li et al., 2021; Liu et al., 2021b; Stukalov et al., 2021; Min et al., 2023; Zhou et al., 2023; Yang et al., 2024) and RNAs (Nabeel-Shah et al., 2022; Wang et al., 2022), influencing cellular processes. In particular, N protein is able to attenuate protein synthesizing machinery, mRNA binding and cell proliferation (Finkel et al., 2021; Pan et al., 2023). Some data indicate that the ability to influence host cell processes depends, at least partially, on properties of N protein to induce cytoplasmic condensate formation via liquid-liquid phase separation (LLPS). Formation of biomolecular condensates by the N protein of SARS-CoV-2 has been actively studied since the beginning of the pandemic and has been shown in many *in vitro* (Chen et al., 2020; Perdikari et al., 2020; Cubuk et al., 2021; Wang et al., 2021b; Dang et al., 2023; Stuwe et al., 2024; Kathe et al., 2024) and *in vivo* research (Iserman et al., 2020; Jack et al., 2021; Lu et al., 2021; Wang et al., 2021b; Wu et al., 2021; Huang et al., 2024; Zhao et al., 2021). SARS-CoV-2 N protein condensates were attributed the functions of accumulating viral replicative apparatus (Savastano et al., 2020) and suppressing host cell response by sequestering proteins involved in signal transduction (Wu et al., 2021). However, some data indicate that interaction with viral genomic RNA can inhibit LLPS (Iserman et al., 2020).

Viral infection can cause the formation of cellular biomolecular condensates, namely stress granules (SGs). The formation of SGs represents a cellular defense mechanism that promotes stress-induced translational arrest through the accumulation of untranslated messenger ribonucleoproteins (mRNPs). The SG response is associated with antiviral innate immunity, and different viruses have evolved various strategies to hijack it (Li and Wang, 2023). The interaction between viral N proteins and cellular proteins that form SGs has been demonstrated in numerous studies (Gordon et al., 2020; Li et al., 2022; Lu et al., 2021; Nabeel-Shah et al., 2022; He et al., 2023; Murigneux et al., 2024; Yang et al., 2024), including those on the lungs of infected patients and mice (Liu et al., 2022). This interaction has been shown to inhibit SG formation, thereby promoting viral infection. The consensus regarding the nature of the formed condensates is yet to be reached, with some data indicating that the SARS-CoV-2 N protein may remodel SGs into atypical foci with an altered protein composition, thereby suppressing innate immunity (Gordon et al., 2020; Li et al., 2022; Lu et al., 2021; Nabeel-Shah et al., 2022; He et al., 2023; Murigneux et al., 2024; Yang et al., 2024). This may vary depending on the type of cells involved (Kim et al., 2022).

It can be reasonably deduced that SARS-CoV-2 N protein may exert an influence on the induction of SGs, and that these properties of this protein may prove to be relevant to the overall pathogenicity of the virus. The potential effects of N proteins from different HCoVs on SG formation have yet to be investigated. In this study, we compare the capacity of all HCoV N proteins to induce cytoplasmic condensates during transient expression in cultured human cells. Our findings indicate that the N proteins of SARS-CoV and SARS-CoV-2 exhibited the lowest capacity to induce condensate formation among all other HCoVs. The formation of condensates was not a consequence of overexpression of N proteins, as is typical for proteins that induce liquid-liquid phase separation (LLPS). Rather, the N proteins were integrated into the forming condensates. The ultrastructural organization and content of the condensates were found to be similar to those observed in SGs, suggesting that N proteins of all HCoVs except SARS-CoV and SARS-CoV-2 induce a stress response that results in the formation of SGs.

## Results

### Expression of nucleocapsid proteins leads to the temporary formation of cytoplasmic condensates

According to the published data, the ectopic expression of SARS-CoV-2 N protein results in the formation of cytoplasmic condensates that exhibit a high degree of accumulation of the expressed N protein (Iserman et al., 2020; Wang et al., 2021a; Wu et al., 2021; Huang et al., 2024). Here, we examine the induction of cytoplasmic condensates formed by N proteins fused with EGFP of low and highly pathogenic HCoVs. To keep it concise, the virus name is used in place of the full designation, e.g., EGFP-N^HCoV-229E^ is referred to as N^229E^. Initially, the formation of N protein-containing condensates was examined in a range of human cell lines. The formation of N-positive condensates was observed to occur in a significantly lower proportion of cells in the case of HT1080, U2OS, and A549 compared to HeLa cells, with the highest proportion observed in cells expressing N^MERS^. For example, N^MERS^-containing cytoplasmic condensates were observed in approximately 37% of HeLa cells, 8% of HT1080 cells, 9% of U2OS cells, and 6% of A549 cells (data from a single representative experiment), and therefore, HeLa cells were selected for all subsequent experiments. It should be noted that the percentage of cells with condensates was not consistent from experiment to experiment.

While the ratios and trends remained stable, there was some variability in the absolute number of cells with condensates. Therefore, although we conducted 2-3 independent experiments in all cases, we have only presented the results of one, which we consider to be the most representative.

Figure 1A illustrates the morphological variants of intracellular distributions of N protein distributions at 24 hours post-transfection. In the majority of cells, the ectopically expressed N proteins were observed to be diffusely distributed throughout the cytoplasm. However, in some cells, N proteins were found to accumulate in cytoplasmic condensates (Figure 1A). The size and number of condensates per cell exhibited considerable variation. In all cases, the majority of the expressed N protein was found to be accumulated within the condensate, with a low concentration detected in the cytosol. In cells exhibiting the highest levels of N^SARS-1^ and N^SARS-2^ expression, as determined by fluorescence intensity, the expressed N proteins formed extensive clumps that occupied a significant portion of the cytosol volume (Figure 1B).

**Figure 1.**
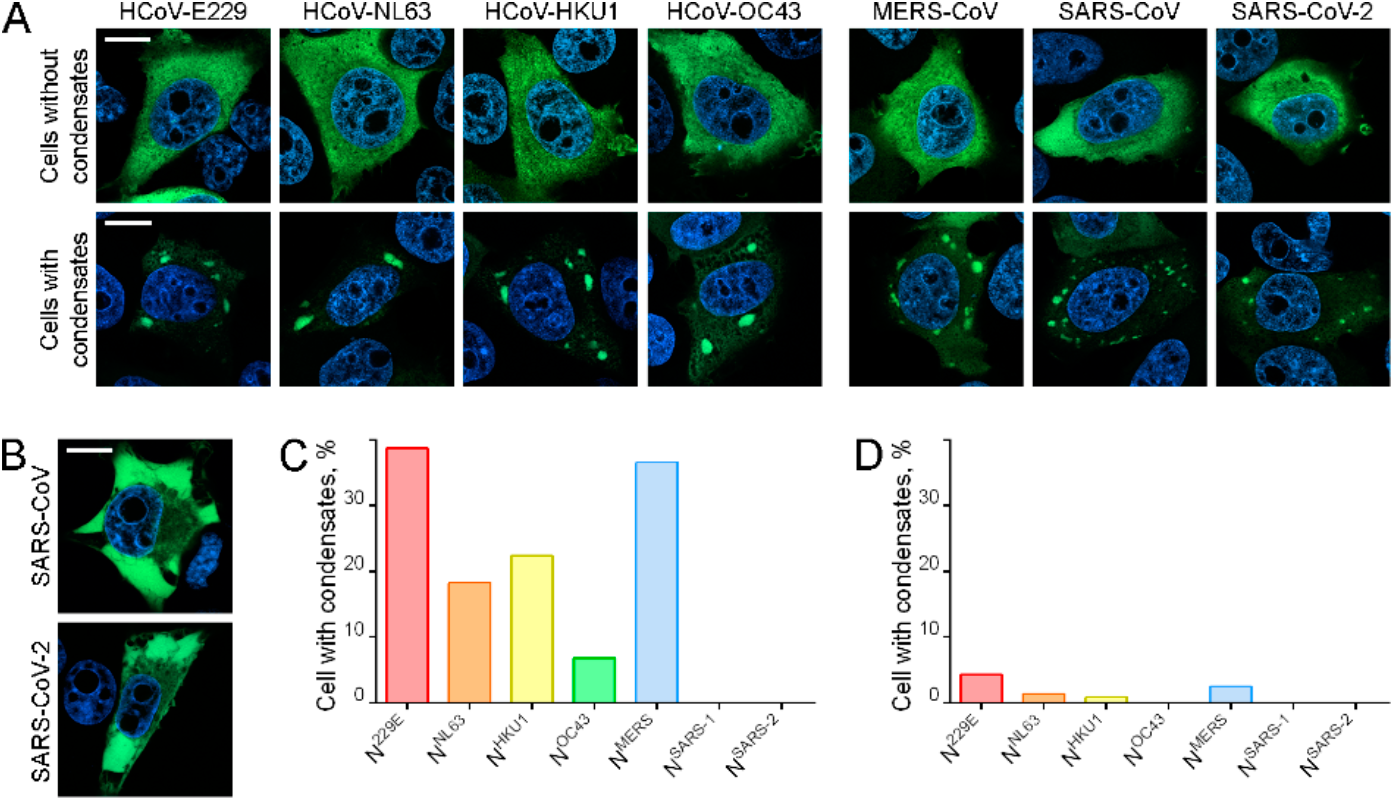
The transient expression of N proteins derived from distinct human coronaviruses resulted in the induction of cytoplasmic biomolecular condensate formation. (**A**) Representative images of N protein-expressing cells without (upper panels) and with (lower panels) cytoplasmic condensates. (**B**) Cells with large condensates (or aggregates) of N protein following the overexpression of N^SARS-1^ and N^SARS-2^. The N protein-containing complexes occupy the majority of the cytoplasm, with the exception of areas in close proximity to the nucleus. (**C**) The percentage of cells with cytoplasmic condensates among all cells expressing N proteins of different HCoVs 24 hours after transfection. (**D**) The reduction in the proportion of cells with N protein-containing cytoplasmic condensates 48 hours after transfection. Scale bars = 10 μm.

The proportion of cells containing condensates exhibited variability among the N proteins of various viruses at 24 hours post-transfection (Figure 1C). A notable proportion of cells (exceeding 10%) exhibited the formation of condensates in the case of the expression of N^229E^, N^NL63^, N^HKU1^, and N^MERS^. The proportion of cells with cytoplasmic condensates never exceeded 2% in the case of N^SARS-1^ and N^SARS-2^, and in some experiments, no cells with condensates could be detected. N^OC43^ exhibited an intermediate position, with a proportion of cells with condensates that did not exceed 10%.

The proportion of cells containing condensates decreased significantly at 48 hours post-transfection, indicating that N protein-containing cytoplasmic condensates were only transient structures (Figure 1D). To determine the potential mechanisms underlying the disappearance of cells with cytoplasmic condensates, we conducted long-term live-cell imaging in 4D mode, i.e., we acquired full stacks of optical sections of each selected cell at one-hour intervals over a period of 16 hours. In all experiments, cells without condensates, which served as controls, were imaged concurrently with cells exhibiting condensates. During the course of the live-cell imaging, a few potential outcomes for cells containing N-positive condensates were observed (Figures 2A-D). Some cells underwent apoptosis within the initial observation period of several hours (Figures 2A), while others retained their morphological integrity and granule content for the entire duration of the experiment, which spanned 16 hours. (Figure 2B). Additionally, the dissolution of condensates was observed in some cells (Figure 2C). On occasion, following the dissolution of condensates, cells were observed to undergo division (however, mitosis of cells with condensates has never been observed) (Figure 2D).

**Figure 2.**
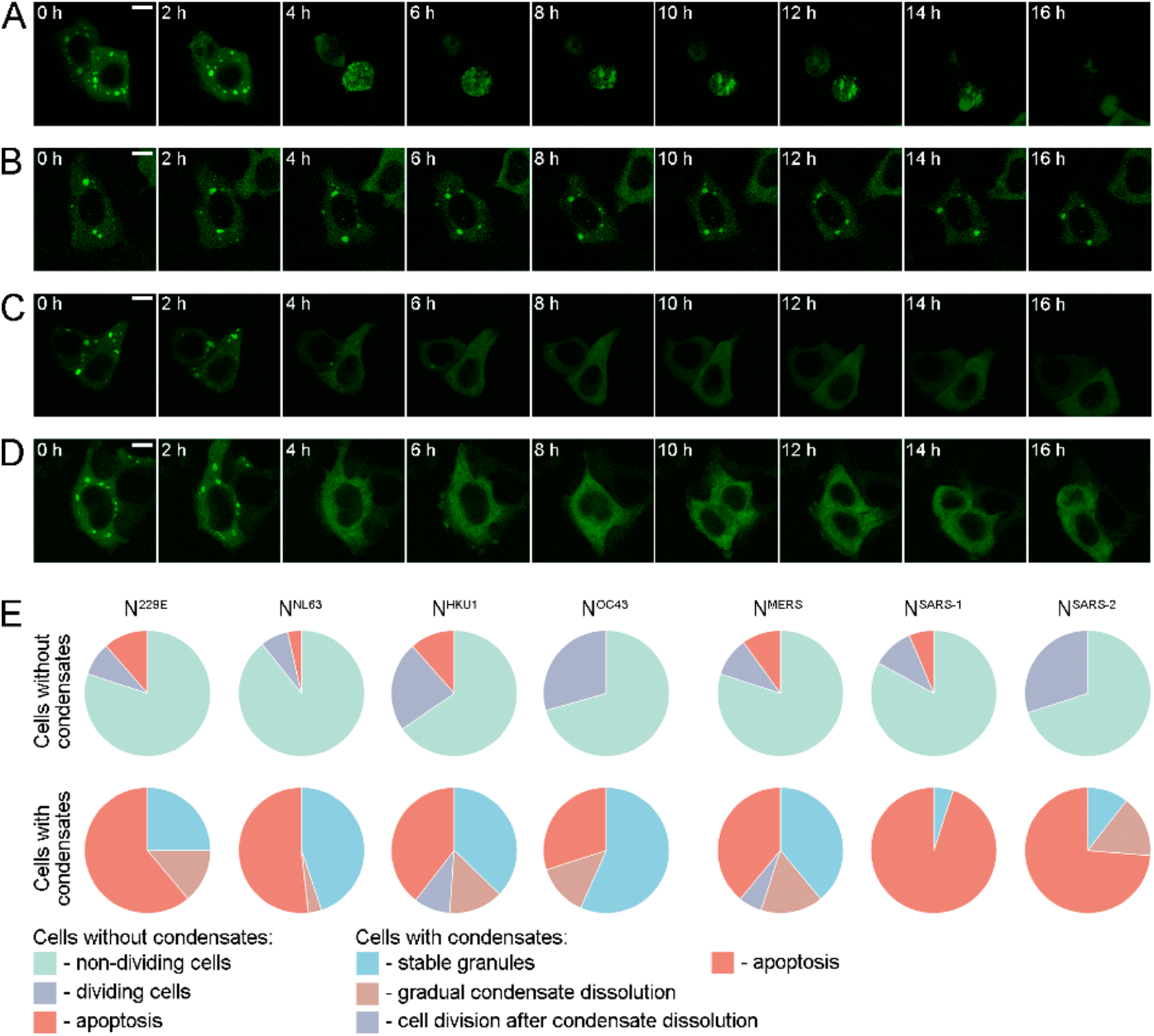
Processes leading to the disappearance of cells with N protein containing condensates. (**A**) A living cell expressing N^MERS^ is undergoing apoptosis during the 16-hour image acquisition period. (**B**) A cell expressing N^MERS^ demonstrates the persistence of condensates throughout the duration of the live image. (**C**) The N^MERS^-containing condensates gradually dissolve. (**D**) A cell expressing N^MERS^ undergoes mitosis after the dissolution of condensates (this mitosis occurred between eight and ten hours). Similar outcomes were observed following the expression of the N proteins of other HCoVs. (**E**) Pie charts represent the events observed during the 16-hour live-cell imaging of N protein-expressing cells with or without condensates. Cells without condensates either remained unchanged throughout the imaging, divided, or died by apoptosis. In cells with condensates, either a gradual dissolution of the condensates occurred (after which the cell could even enter mitosis, i.e. its condition returned to normal), or the cells died by apoptosis. Scale bars = 10 μm.

To ascertain the frequencies of the various outcomes observed in cells containing condensates, we acquired images of at least 20 cells with condensates and an equivalent number of control cells without condensates for each N protein (Figure 2E). A relatively small percentage of cells without condensates (up to 11%) underwent apoptosis, while the majority of these cells remained unchanged throughout the experiment. Some of the control cells initiated a cell division (mitosis), indicating that the live cell imaging did not significantly affect the cells (since under conditions of cellular stress, progression through the cell cycle is usually inhibited). A significant proportion of cells with condensates underwent apoptosis, indicating a reduction in cell survival compared to cells without condensates. An extreme case was observed in cells with expression of N^SARS-1^ and N^SARS-2^, which exhibited a particularly high fraction of cellular death events (95% for N^SARS-1^ and 73% for N^SARS-2^, respectively). As previously stated, the N proteins of these two closely related viruses exhibited a markedly low prevalence of cells with condensates. Thus, the disappearance of cells with N protein-containing cytoplasmic condensates was the result of two processes: cell death and normalization of cell morphology via condensate dissolution.

### N protein overexpression did not not induce condensate formation

The precise mechanism of N protein-containing cytoplasmic condensate formation *in vivo* remains unclear. Some data indicate that the formation of SARS-CoV-2 N protein condensates is associated with the overexpression of this protein, i.e. an increase in protein concentration can result in the induction of the formation of membraneless structures via the LLPS mechanism (Iserman et al., 2020). In contrast, recent data suggest that elevated levels of N protein may reduce the propensity for cytoplasmic condensate formation (Yang et al., 2024). To investigate the role of N protein concentration in the biogenesis of cytoplasmic condensates, we measured the total EGFP fluorescence of cells with and without cytoplasmic condensates (due to the scarcity of cells with condensates, we were unable to conduct such analysis for N^SARS-1^ and N^SARS-2^).

Total fluorescence of cells with and without condensates was measured 24 hours after transfection (Figure 3A). No increase in the amount of protein was observed in cells with condensates compared to cells without condensates. Conversely, in most cases, the median was slightly lower, and among the cells with overexpression, there were no cells with condensates. It is noteworthy that the expression level of N^OC43^ (less than 10% of cells with condensates) and especially of N^SARS-1^ and N^SARS-2^ (single cells with condensates) was higher than that observed in the case of other proteins. In the case of N^OC43^, the brightness of cells with condensates was found to be significantly lower in comparison to cells without condensates. This observation is consistent with the data that overexpression of N proteins can suppress condensate formation (Yang et al., 2024).

**Figure 3.**
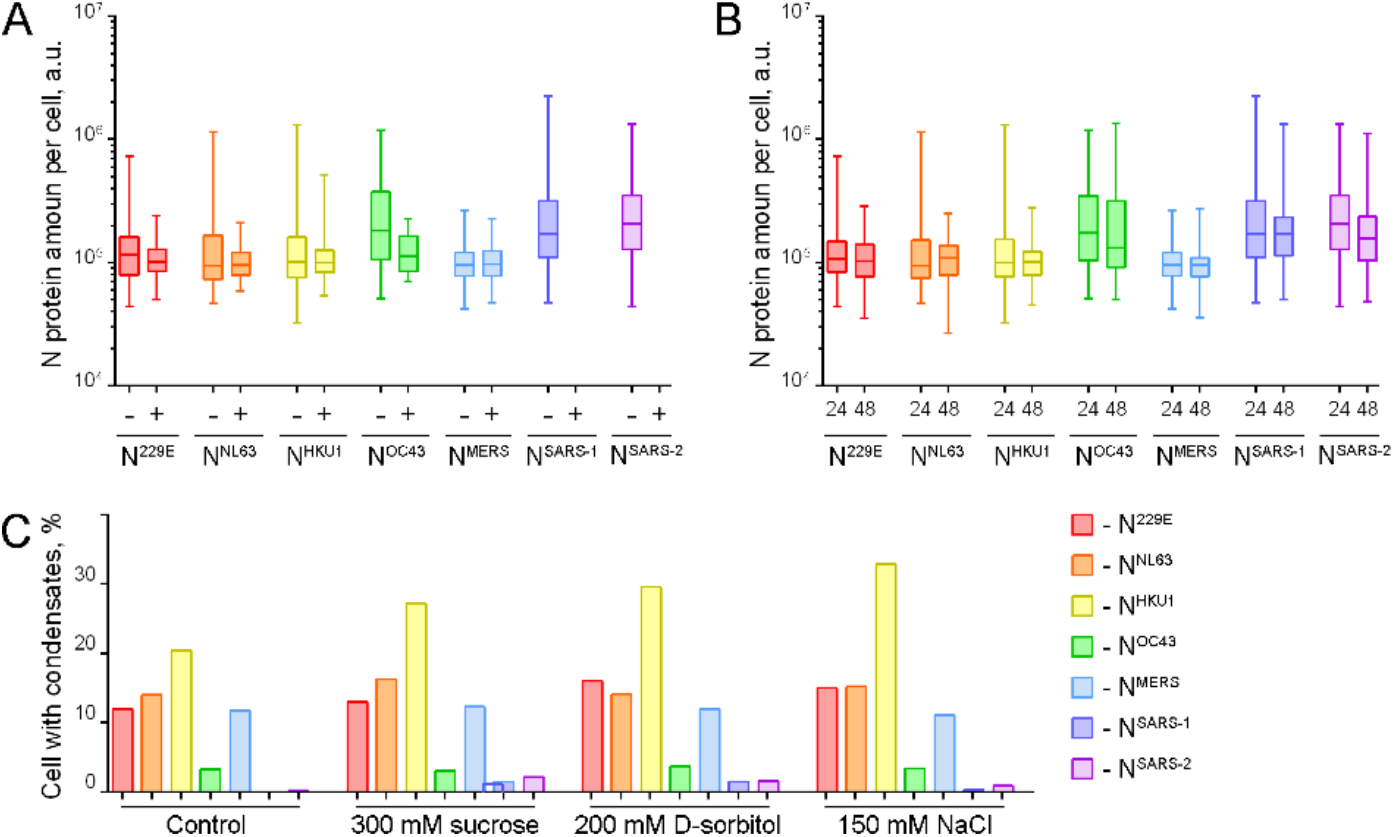
Comparative analysis of protein amount among HCoVs N proteins in cell. (**A**) Total fluorescence measurements in N protein-containing cells without condensates (-) and with condensates (+) 24 h after transfection. (**B**) Comparison of N protein amount in cells by measuring total fluorescence 24 h and 48 h after transfection. (**C**) N protein-containing cells were treated with hyperosmotic solutions (300 mM sucrose, 200 mM D-sorbitol, 150 mM NaCl) to induce the formation of condensates by reducing cell volume.

As illustrated in Figures 1C and D, the number of cells containing N protein-containing condensates was reduced at 48 hours post-transfection. If the hypothesis that N protein overexpression promotes condensate formation is correct, one can anticipate a decline in the ectopically expressed N protein amount at 48 hours post-transfection. The protein content of the cells was analyzed at 24 and 48 hours after transfection. In this experiment, EGFP fluorescence was measured in all cells, as at 48 hours post-transfection, cells with condensates were exceedingly rare and it was challenging to measure them separately (Figure 2B). For all HCoVs, the N protein content was not significantly reduced, although the number of cells with condensates was reduced, as shown above (Figures 1C and D).

Finally, an attempt was made to elevate the N protein concentration within the cells through the use of hyperosmotic solutions. It has been demonstrated that the formation of specific condensates is concentration-dependent, and that their formation can be induced by hyperosmotic treatment, which reduces the cell volume (Musinova et al., 2016; Yasuda et al., 2020; Boyd-Shiwarski et al., 2022; Gao et al., 2022). The culture medium was supplemented with sucrose (300 mM), D-sorbitol (200 mM), or NaCl (120 mM) in order to modulate the osmotic conditions. During visual analysis, no significant formation of condensates was observed in the cytoplasm of cells that did not initially contain condensates. This phenomenon was observed for N proteins of all seven HCoVs except N^HKU1^, for which a notable increase in cells with condensates was observed (Figure 3C). However, even in the case of N^HKU1^, only a moderate increase in the proportion of cells with condensates was observed (32.9% for NaCl treatment compared to 20.5% in isotonic medium).

The aforementioned estimates indicate that overexpression (an increase in the amount of N protein per cell) did not induce cytoplasmic condensate formation via LLPS in vivo. Instead, it is probable that this process was inhibited, as was observed in the case of the N^OC43^ and, in particular, the N^SARS-1^ and N^SARS-2^ proteins.

### The ectopically expressed HCoV N proteins were integrated into SGs

A number of published papers indicate that the N protein of SARS-CoV-2 can accumulate within SGs induced by treatment with polyU RNA or sodium arsenite. According to the data presented here, ectopically expressed N proteins are integrated into cytoplasmic condensates, which can be assumed to be SGs. In order to evaluate this claim, we analyzed some features of the N protein-containing condensates.

The formation of SGs is a consequence of translation inhibition (Marcelo et al., 2021). To visualize protein biosynthesis in cells with biomolecular condensates, we employed a click chemistry-based approach to detect the newly synthesized proteins. Our observations revealed that 100% of cells with N protein-containing condensates exhibited complete translational arrest. As a control, we utilized neighboring cells with N protein expression but without condensates, in which translation was never inhibited (Figure 4).

**Figure 4.**
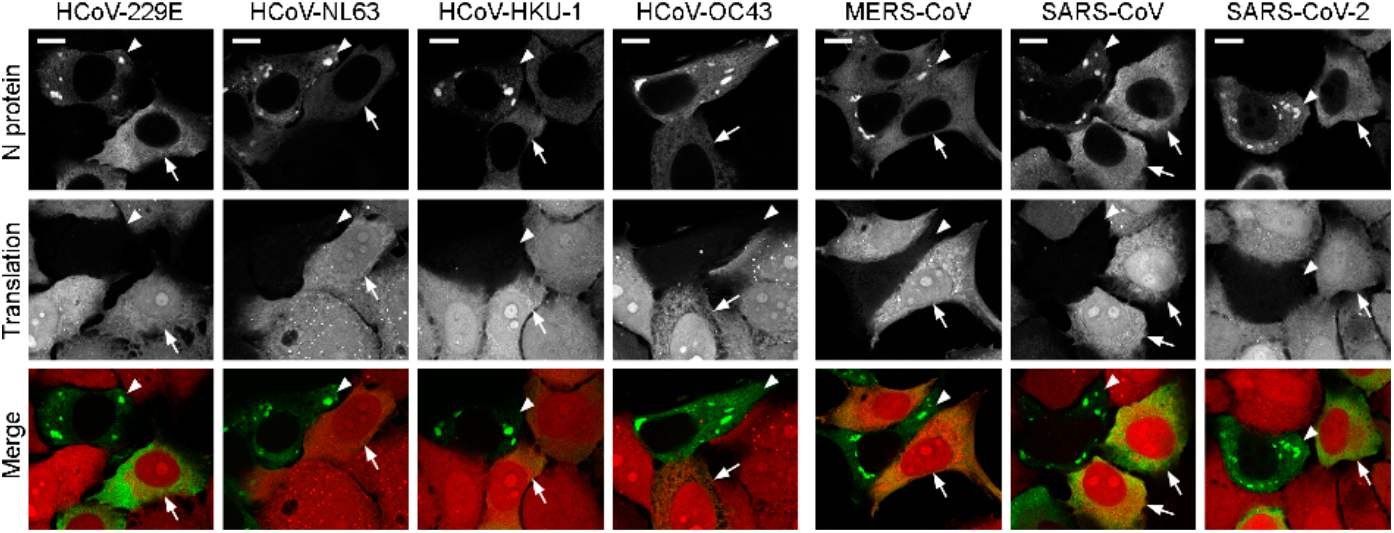
Translation inhibition of cells with N protein-containing condensates. Cells with condensates are marked with arrowheads, control cells without condensates are marked with arrows. N protein - green, click-labeled translation products - red) Scale bars = 10 μm.

The published data indicate that SGs accumulate messenger RNA (mRNA) and small ribosomal subunits (Reineke et al., 2012). In all cases involving seven HCoV N proteins, the accumulation of poly-A RNAs was clearly observed. Moreover, N-containing granules were found to contain components of the small ribosomal subunit, specifically 18S ribosomal rRNA (Figure 5). It is noteworthy that while the majority of poly-A-containing RNA molecules were observed to accumulate within the condensates, only a limited number of 18S rRNA molecules were seen inside the condensates, with the majority of molecules still distributed throughout the cytoplasm. However, the detection of other types of eukaryotic rRNAs, specifically 5.8S rRNA and 28S rRNA, demonstrated that the condensates were depleted of large ribosomal subunits (Figure 5).

**Figure 5.**
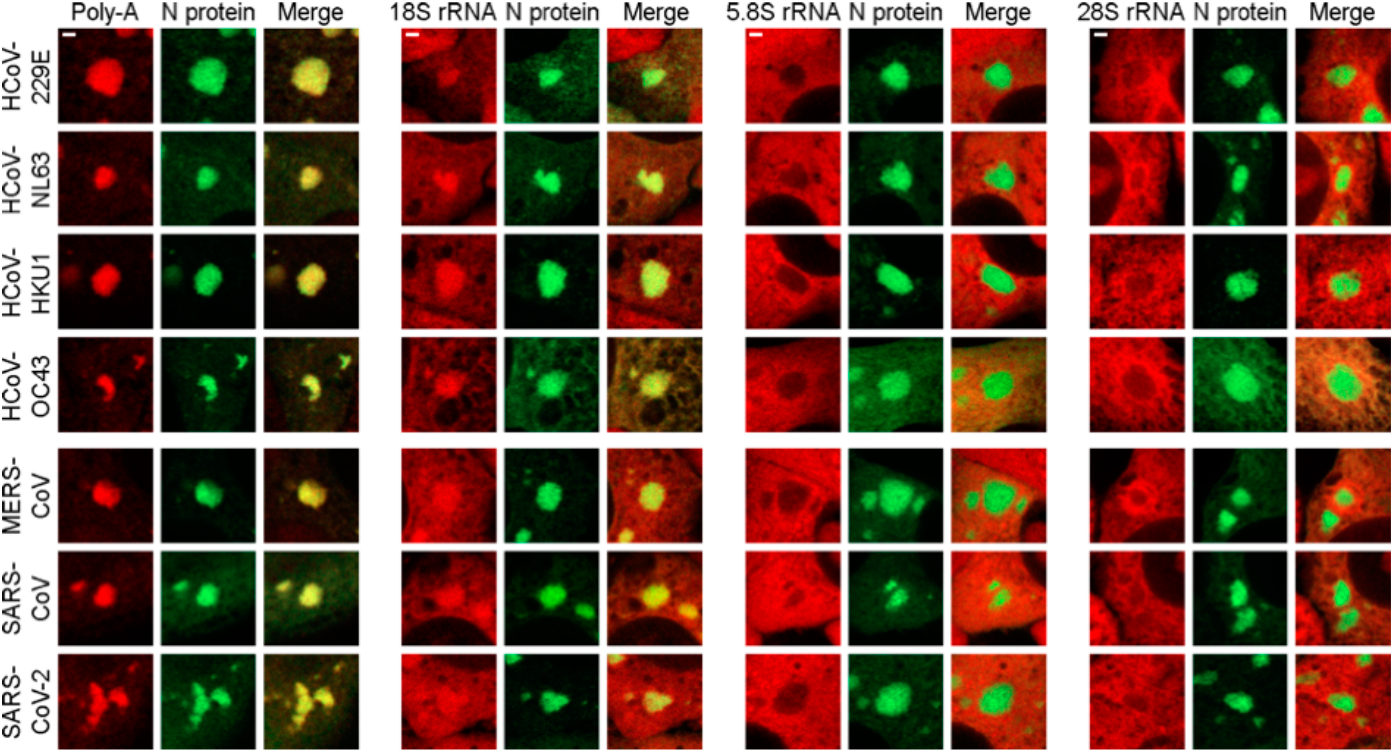
Detection of RNAs accumulated in N protein-containing condensates by fluorescence *in situ* hybridization (FISH). Poly-A and 18S rRNA accumulate within the condensates, while 5.8S and 28S rRNAs are preferentially distributed outside the condensates. Scale bars = 1 μm.

**Figure 6.**
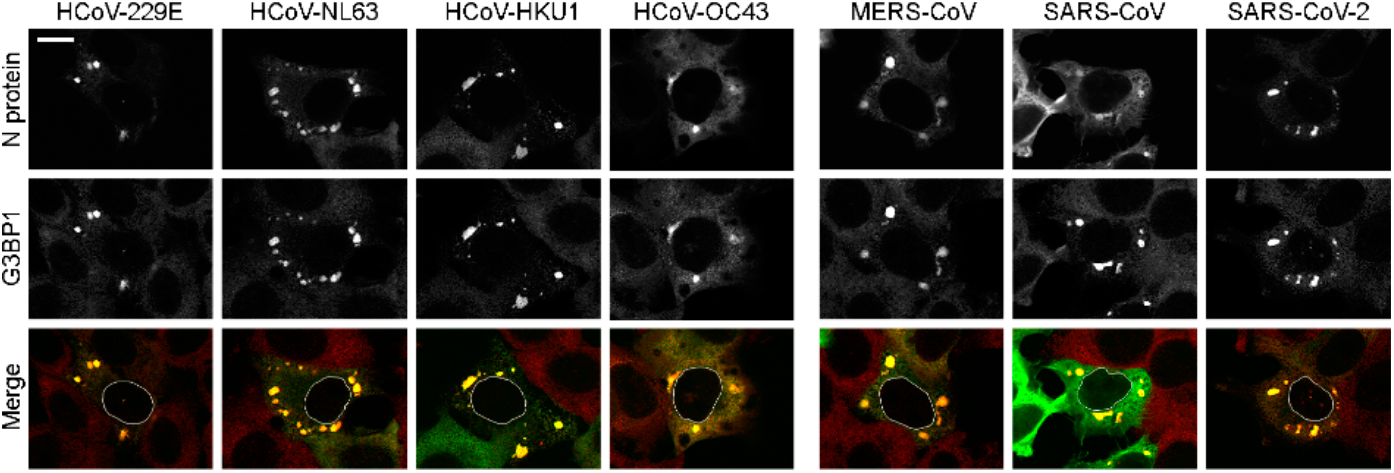
Accumulation of SG marker G3BP1 protein within N-protein-containing condensates. In the merge images (N proteins - green, G3BP1 - red), the nuclear contours of cells with condensates were added. Scale bar = 10 μm.

To further confirm the similarity of the N-positive condensates to SGs, we stained cells with antibodies against one of the major components of SGs - G3BP1, the protein that induces phase separation leading to the formation of stress granules [42]. We observed strong colocalization of all seven N protein-containing condensates with G3BP1 (Figure 7).

**Figure 7.**
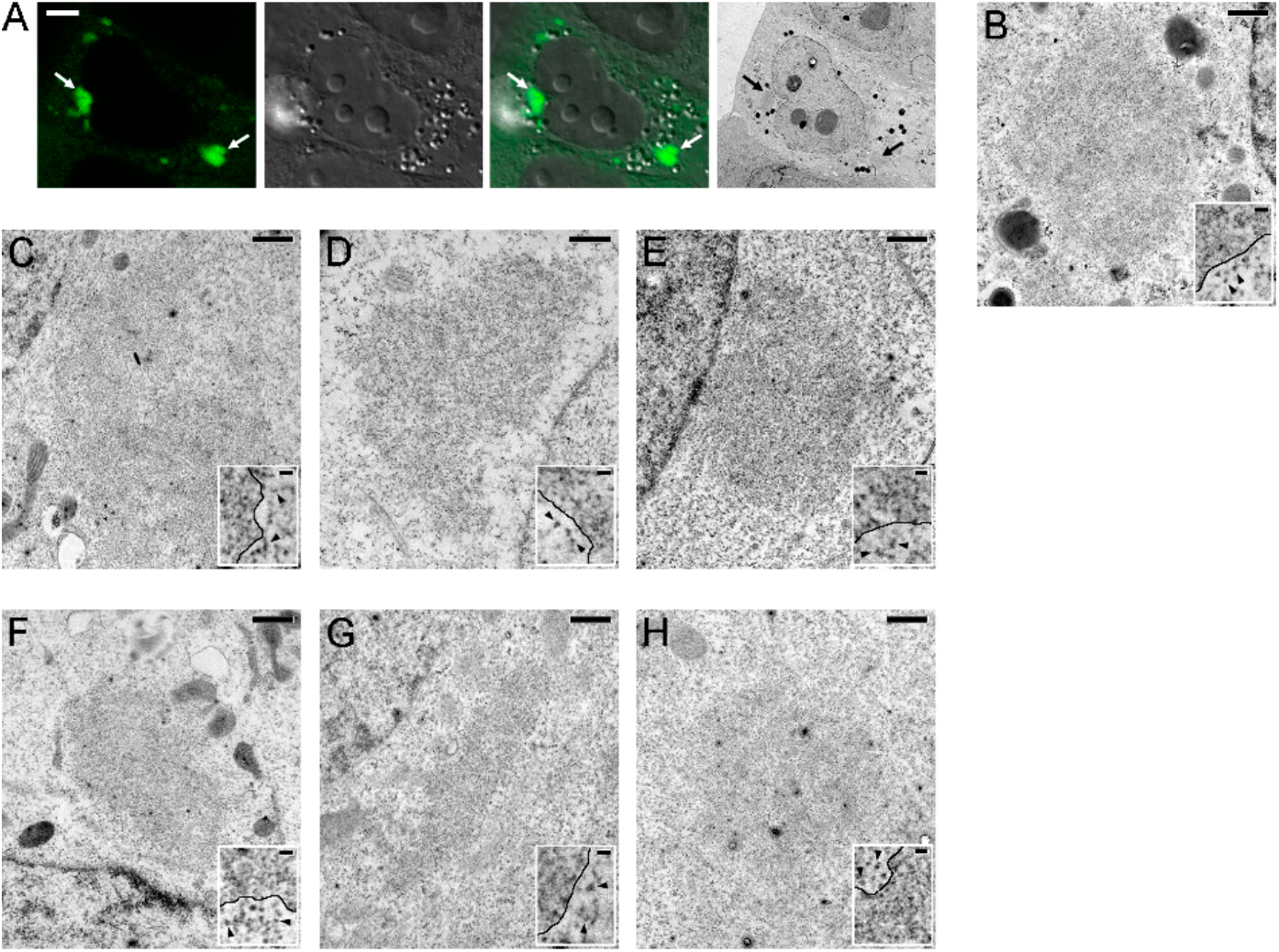
Ultrastructural organization of cytoplasmic N protein-containing condensates. (**A**) correlative light and electron microscopy (CLEM) of N^229E^-protein-containing condensates. The fluorescence, Differential interference contrast (DIC), and transmission electron microscopy images were acquired, allowing the identification of N protein-containing condensates (arrows) in ultrathin sections without specific staining. It is noteworthy that the cytoplasm of the cell with condensates is lighter than the cytoplasm of the surrounding cells, which may be associated with the condensation of a significant amount of ribosomal material in the granules. (**B-H**) Electron microscopy images of condensates containing N^229E^ (**B**), N^NL63^ (**C**), N^HKU^1 (**D**), N^OC43^ (**E**), N^MERS^ (**F**), N^SARS-1^ (**G**), N^SARS-2^ (**H**). Inserts illustrate the organization of the condensate periphery. The boundaries of the condensates are indicated by black lines, the ribosomal particles are indicated with arrowheads. Scale bar = 5 μm (A), 0.5 μm (B-H), 50 nm (inserts in B-H).

Then, we analyzed the ultrastructural organization of N protein-containing cytoplasmic condensates of all seven HCoVs. Due to the rarity of such cells, especially in the case of N^SARS-1^ and N^SARS-2^, we performed the study using correlative light and electron microscopy (CLEM). The combination of light and electron microscopic images allowed accurate identification of cells with condensates (Figure 7A). The cytoplasm of cells with condensates was significantly lighter compared to normal cells (Figure 7A). For all seven N proteins, cytoplasmic condensates appeared as membraneless structures (Figure 7B-H). Analysis of high magnification images showed that the condensates were composed of finely fibrillar material arranged in a dense network. This material was easily distinguished from ribosomes (or, probably, large subunits of ribosomes), discrete bodies that were clearly visible in the cytoplasm (insets in Figure 7B-H).

## Discussion

HCoV N proteins are highly abundant in late stages of viral infection and are not only important for virus particle formation, but can also interact with cellular components and influence cellular responses. Currently, the properties of N proteins are poorly understood and researchers are mainly focused on highly pathogenic coronaviruses (especially, SARS-CoV-2). Comparison of N proteins of different - low and highly pathogenic - HCoVs may help to elucidate possible mechanisms of disease progression and identify potential targets for medical intervention. Here, we performed a comparative analysis of the ability of N proteins from all human coronaviruses to induce the formation of biomolecular condensates during transient expression in human cells. We analyzed the N proteins of all human coronaviruses, including highly pathogenic (MERS-CoV, SARS-CoV, and SARS-CoV-2) and low pathogenic viruses (HCoV-229E, HCoV-NL63, HCoV-HKU1, and HCoV-OC43). Expression of N proteins led to the formation of N protein-containing condensates in the cytoplasm. N proteins N^229E^, N^HKU-1^, N^NL63^ and N^MERS^ were most prone to form condensates, while N^SARS-1^ and N^SARS-2^ were characterized by insignificant condensate formation. N^OC43^ showed intermediate properties in the majority of experiments.

The formation of N protein-containing cytoplasmic condensates was observed to be transient, with the majority of these structures disappearing by 48 hours. Live cell imaging demonstrated that the condensates either dissolved or that cells with condensates underwent death via apoptosis. It is worth noting that these experiments showed that N^SARS-1^ and N^SARS-2^ differed from other HCoVs. In N^SARS-1^ and N^SARS-2^, cells exhibited a proclivity for undergoing apoptosis, whereas in all other HCoVs, the dissolution of condensates was markedly more prevalent. It appears that N^SARS-1^ and N^SARS-2^ have an inhibitory effect on the formation of condensates, and the formation of condensates in the case of these proteins was observed only in instances where apoptosis was the preferred outcome.

The data presented in this study demonstrate two characteristics associated with the formation of condensates containing N proteins. Firstly, the formation of these condensates is not contingent on the overexpression of N proteins. Conversely, in all instances, cells exhibiting N protein overexpression did not exhibit the formation of N protein-containing condensates. It would appear that overexpression has the effect of inhibiting the formation of condensates. These observations are in accordance with the findings of Yang et al. (Yang et al., 2024) and in disagreement with the findings of Iserman et al. (Iserman et al., 2020).

Secondly, our findings demonstrated that N protein-containing condensates exhibit a striking resemblance in composition to SGs. The formation of these membraneless organelles occurred in cells with suppressed translation, and the condensates themselves exhibited a composition similar to that of typical SGs. Specifically, poly-A RNA and rRNA of the small 40S subunit (18S rRNA) were observed to accumulate in the condensates, whereas rRNAs of the large 60S subunit (5.8 and 28 S rRNA) were not. Furthermore, the proteins that are crucial for SG formation - G3BP1 - were observed to accumulate in these condensates. In addition, we found that N protein-containing condensates had an ultrastructure very similar to that of SG obtained by arsenite treatment (Souquere et al., 2009). Therefore, it appears that N protein expression induces stress that ultimately leads to SG formation, and subsequently, N proteins are incorporated into these SGs.

Thus, N proteins of different HCoVs do not induce condensate formation by LLPS alone after transient expression in cultured human cells. Rather, they induce cellular stress, which results in the inhibition of translation and the formation of SGs by LLPS. Consequently, N proteins appear to be incorporated into these SGs. Notably, two N proteins from closely related viruses, SARS-CoV and SARS-CoV-2, appear to exhibit either the lowest ability to induce SG formation or the highest ability to antagonize SG formation. These observations are in accordance with the data indicating that the SARS-CoV-2 N protein antagonizes SG assembly (Li et al., 2022; Liu et al., 2022; Yang et al., 2024). However, a noteworthy novel observation is that this activity may be exclusive to only two highly pathogenic HCoVs - SARS-CoV and SARS-CoV-2. In light of the limitations of the experimental model employed, further research utilizing more sophisticated experimental models is required to elucidate the specific properties of individual proteins belonging to distinct HCoV species.

## Materials and methods

### Cell culture

Cultures of human carcinoma cells (HeLa), fibrosarcoma cells (HT1080), osteosarcoma cells (U2OS), and alveolar basal epithelial cells (A549) were cultivated in DMEM medium (Paneco) supplemented with 10% heat-inactivated fetal bovine serum (Gibco), 2 mM alanine-glutamine (Paneco), and an antibiotic-antimycotic solution (Gibco). Cell transfection was conducted using either Lipofectamine 2000 reagent (Invitrogen, Cat. No. 11668019) or GenJect-40 (Molecta, Cat. No. Gen40-1000p) according to the manufacturers’ instructions.

### Plasmid construction

The following plasmids were used for plasmid construction: for SARS-CoV-2 pGBW-m4134905 was a gift from Ginkgo Bioworks & Benjie Chen (Addgene plasmid # 151966), for HCoV-OC43 pGBW-m4134899 was a gift from Ginkgo Bioworks & Benjie Chen (Addgene plasmid # 151902), for HCoV-NL63 pGBW-m4134908 was a gift from Ginkgo Bioworks & Benjie Chen (Addgene plasmid # 151957), for HCoV-HKU1 pGBW-m4134897 was a gift from Ginkgo Bioworks & Benjie Chen (Addgene plasmid # 151948) (Addgene plasmid, #151948), for HCoV-229E pGBW-m4134909 was a gift from Ginkgo Bioworks & Benjie Chen (Addgene plasmid # 151901). Sequences of nucleocapsid proteins of SARS-CoV and MERS-CoV were obtained from the NCBI database (NC_004718.3 for SARS-CoV and JX869059.2 for MERS-CoV) and synthesized (Evrogen, Russia). Full-length nucleocapsid protein genes were amplified by PCR using Phusion high fidelity DNA polymerase (Thermo Scientific). The amplified PCR products were digested with restriction enzymes and inserted into EGFP-C1 vector (Clontech).

### Hypertonic stress

To induce the formation of biomolecular condensates, HeLa cells transfected with plasmids encoding nucleocapsid proteins were treated with sucrose (Sigma) at a concentration of 300 mM, D-sorbitol (Sigma) at a concentration of 200 mM, and NaCl (Fluka) at a concentration of 120 mM (solutions were prepared in full culture medium), then cells were fixed in 3.7% paraformaldehyde, permeabilized with 0.5% Triton in PBS, washed with 1xPBS, stained with DAPI.

### Fluorescent microscopy

For live-cell imaging cells were grown in 35-mm coverslip dishes. Imaging was performed on a Nikon C2 confocal microscope with a Plan Apo 60x (NA 1.4) objective at 37°C and focus stabilization using a PFS system (Nikon). Temperature, CO_2_, and humidity was maintained using Stage Top Chamber H301-K-FRAME (Okolab). A laser with a wavelength of 488 nm was used to excite EGFP. Cells were recorded in automatic recording mode at a frequency of 1 time per hour for 16 hours.

For fluorescence intensity estimations, preparations were photographed using Axiovert 200M microscope (Carl Zeiss) equipped with Plan Neofluar 20x objective. Image processing was carried out in the image processing package Fiji (https://imagej.net/software/fiji/).

### Translation

Protein synthesis was detected using the Click-iT™ Plus OPP Alexa Fluor™ 594 Protein Synthesis Assay Kit (Thermo, Cat. No. C10457), according to the manufacturer’s instructions.

### Fluorescence *in situ* hybridisation (FISH)

Cells were fixed with 3.7% paraformaldehyde for 10 min, then fixed in 100% ice-cold methanol for 10 min, then 70% ethanol was added for 10 min. The cells were then incubated in a hybridization buffer (yeast tRNA 1 mg/ml, BSA 0.005%, dextran sulfate 10%, formamide 25%, 2xSSC) with 18S rRNA (5′-GATCAACCAGGTAGGTAAGGTAGAGCGCGGCGA-cy3), 28S rRNA (5′-CGTGCGTGCGCGCGCGCCTGTTTGGGAAC-cy3) (Carron et al., 2012), 5.8S rRNA (5’-TCCTGCAATTCACATTAATTCTCGAGCTAGC-cy3) (Pietras et al., 2022), or Oligo-dT 25 probe at a concentration of 1 μg/ml overnight at 40°C C in a humid chamber. Samples were washed in 4xSSC, then in 2xSSC, stained with DAPI and mounted in Mowiol 4-88 (Aldrich, Cat. No. 81381) containing the anti-bleaching agent DABCO (Sigma, Cat. No. D-2522).

### Immunocytochemistry

Cells were fixed with 3.7% paraformaldehyde for 15 min, washed with 1xPBS, permeabilized with 0,5% Triton for 10 min and stained with antibodies against G3BP1 (Santa Cruz Biotechnology, Cat. No. sc-365338) for 1h. For secondary antibodies, Donkey Anti-Rabbit IgG AlexaFluor 568 (Abcam, Cat. No. Ab175692) were used.

### Confocal microscopy

For morphology analysis of fixed cells, we used the LSM900 confocal microscope (Carl Zeiss) (provided by the Moscow State University Development Program). The final images were processed using ImageJ for brightness/contrast correction (excepting Figure 4, where gamma correction was also performed to better visualize cytoplasm boundaries).

### Correlative light and electron microscopy (CLEM)

For CLEM experiments, cells were grown in 35 mm coverslip dishes and prefixed in 3.7% formaldehyde with 0.1% glutaraldehyde in 0.1 M cacodylate buffer for 15 minutes. Cells with condensates were imaged by confocal microscopy in both EGFP and DIC channels. To facilitate identification of the imaged cells after embedding, we outlined them with a syringe needle. Subsequently, an additional fixation was performed in 2.5% glutaraldehyde in 0.1 M cacodylate buffer for 8 hours, followed by a 1-hour postfixation with 1% osmium tetroxide.

The samples were then dehydrated in ethanol and acetone (70% ethanol contained 2% uranyl acetate), and finally embedded in Spi-pon 812 (SPI, Cat. No. 02660-AB). After embedding, cells of interest were identified by phase-contrast microscopy. Ultrathin sections were prepared on an Ultracut E ultratome (Reichert Jung). Sections were stained with lead citrate and photographed using a JEM-1400 electron microscope (Jeol).

## Acknowledgments

We thank Ginkgo Bioworks & Benjie Chen for a generous gift of plasmids, and A. Lasarev for invaluable assistance with electron microscopy.

